# MurSS: Multi-resolution Selective Segmentation Model for Breast Cancer

**DOI:** 10.1101/2023.10.10.561807

**Authors:** Joonho Lee, Geongyu Lee, Tae-Young Kwak, Sunwoo Kim, Min-Sun Jin, Chungyeul Kim, Hyeyoon Chang

## Abstract

We propose the Multi-resolution Selective Segmentation model (MurSS) for segmenting benign, Ductal Carcinoma In Situ, and Invasive Ductal Carcinoma in breast resection Hematoxylin and Eosin stained Whole Slide Images. MurSS simultaneously trains on context information from a wide area at low resolution and content information from a local area at high resolution, aiming for a more accurate diagnosis. Additionally, through the selection stage, it provides solutions for ambiguous tissue regions. Our proposed MurSS achieves a mean Intersection of Union performance of 91.1%, which is at least 16.8% and at most 19.0% higher than well-known image segmentation models.

## 1. Introduction

Breast cancer is one of the most commonly diagnosed cancers and is the leading cause of cancer death in women. Therefore, predicting breast cancer prognosis is an essential area of study. In general, Extensive Intraductal Components (EICs) are an important factor in predicting the prognosis of breast cancer [1] [2]. To calculate the EIC, the area of Ductal Carcinoma In Situ (DCIS) and Invasive Ductal Carcinoma (IDC) regions is required [3]. We propose a MurSS to automatically segment DCIS and IDC regions based on deep learning. Pathological images require a lot of computational cost due to their high resolution. For instance, the resolution of Whole Slide Image(WSI)s from The Cancer Genome Atlas (TCGA) dataset [4], widely used in pathological image analysis, is giga pixels *×* giga pixels which is significantly higher than the resolution of standard image datasets. Therefore, each WSI is divided into smaller pieces called patches, and the model is trained using these patches. When extracting a patch at the highest resolution, the wide area’s contextual information is not used because there is available tissue around the patch that is not considered in the receptive field. Also, when extracting a patch at the lowest resolution with the same size, the same receptive field contains more contextual information than before. To utilize such feature information diversely, there are studies that aim to use multi-resolution patches in patch classification tasks or segmentation tasks [5] [6]. Follow these previous studies, we propose MurSS that observes WSIs from both low and high resolutions and combines the two feature information using adaptive normalization to segment benign, DCIS, and IDC more accurately. Furthermore, the morphological similarities between DCIS and IDC can adversely affect the model’s training. In reality, the same type of ductal carcinoma is categorized as either DCIS or IDC based on whether it’s inside the duct [7], and pathologists typically combine observations for accurate diagnosis. Such ambiguous annotations on pathological images can induce data uncertainty, leading to inconsistent diagnostic results. To address these uncertainties, we developed a method named ‘Selective Segmentation’ that evaluates the uncertainty of regions. By utilizing the MurSS, the performance based on mean Intersection of Union(mIoU) increased by 14.1% compared to models that did not use the selective segmentation method. Also, MurSS achieves a mIoU performance of 91.1%, which is at least 16.8% and at most 19.0% higher than well-known image segmentation models. The major contributions of this study are summarized below.

- Multi-resolution adaptive normalization model to combine multi-resolution information.
- Selective segmentation method to address data uncertainty of DCIS and IDC .

The remaining part of this study is organized as follows. We briefly introduce related work in Section 2. In Section 3, we explain the proposed Multi-resolution Adaptive Normalization model (MurAN) and MurSS, respectively. Section 4 represents experimental results, and Section 5 concludes this study.

## 2. Related work

This section briefly introduces previous studies on various image segmentation models used in medical domains or general images. Convolutional Neural Network(CNN)-based segmentation models, multi-resolution segmentation models, and selective models will be introduced.

### 2.1 CNN based segmentation model

CNN is one of the popular techniques in image segmentation tasks. In the early stage of the segmentation model, Fully-Convolutional Networks (FCN) was proposed by [8], to perform segmentation using CNN filters for any images with different sizes. Inspired by FCN, [9] proposed U-Net to successfully segment cellular structures from the background in medical images. U-Net combines low-level features in the high-to-low downsample process into the same-resolution high-level features in the low-to-high upsample process progressively through skip connections. [10] proposed DeepLabV3, which utilized dilated convolution to expand the receptive field, thereby leveraging broader context information. Most segmentation tasks usually compress high resolution images by passing them through CNN layers and then employing an upsampling method. However, [11] proposed High-Resolution Network(HRNet), which maintains high resolution throughout the entire process and progressively adds feature maps of lower resolutions in parallel as the layers deepen, a technique described as

multi-resolution streams in parallel. As a result, HRNet can accurately segment regions even in complex images since features from high resolution and low resolution are combined. Various models are being studied to leverage connections between features of different scales, or expansion of the receptive field to utilize multi-scale feature maps or wider receptive field.

### 2.2 Multi-resolution based segmentation model

There exists many studies to combine features from different resolutions of images. [12] insist that as input resolution gets higher, more details are generated, but see inconsistencies in the scene structure. [13] propose an Image Cascade Network (ICNet) using cascade feature fusion. ICNet is trained by designed networks for each of three different resolutions and combining them. In this way, the important feature information of each resolution is used. [14] proposed a deep multi-magnification network for multi-class breast cancer image segmentation (DMMN) consisting of three U-Nets. They use the method of inputting three images from different resolutions to the encoder of U-Net and concatenating the features. Various studies have been studied to analyze and combine images at different resolutions.

### 2.3 Selective model

Handling statistical uncertainty is an important part of deep learning. In tasks such as autonomous driving and healthcare, even a small error can cause a big problem. To solve this problem, researchers have been working on statistical uncertainty for a long time, such as the reject option applied to Support Vector Machine [15] and the reject option applied to nearest neighbor or k-nearest neighbor [16] [17]. When considering selective prediction in neural networks, the most common method is to find a threshold to reject from a pre-trained model on the entire data and train using that

threshold [18] [19]. This approach involves training on the entire dataset and determining a reject threshold using the outputs that yield ambiguous results. Subsequently, using this rejection threshold, data is excluded during training. A drawback of this method is that it requires two rounds of training. Therefore, recent studies is devising methods that can simultaneously determine the rejection threshold and exclude data based on that threshold during training [20]. In addition, methods using Monte-Carlo dropout to reject uncertain data [21] and using softmax response are still available.

Based on these studies, this study proposes the MurSS, which combines the MurAN and selective segmentation method. MurSS tries to overcome the limitations of existing methods by combining context and content information at various resolutions and effectively performing segmentation for uncertain areas.

## 3. Method

This section describes the key elements of the MurSS. MurSS consists of MurAN, which receives high resolution and low resolution as input and produces output, and a selection stage that selects the output based on uncertainty. MurAN will be introduced at 3.1 and the selective segmentation method will be introduced at 3.2.

### 3.1 Multi-resolution Adaptive Normalization model

The architecture of our proposed model is shown in Figure 1. It processes two inputs of different resolutions, utilizing a shared-weight CNN model, termed as the ‘backbone’, for extracting multi-resolution features. When high resolution input is fed, the model generates mid and high-level features through the backbone. Conversely, for low resolution input, the model obtains context information containing the overall style of the image after passing through the backbone. High features from high resolution pass through the non-local block layer to bring in patch-wise global features. Non-local Block is a method proposed by [22] and follows the structure of Figure 2 and the equation below.

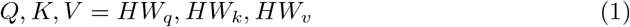

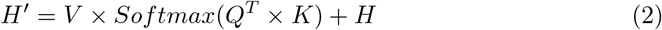

High level features, *H ∈ R*^*C*H*W*^, come out as Query(*Q*), Key(*K*), and Value(*V*) through pointwise convolution (Eq 1), and these values have patch-wise global features*∈*equal to the original feature size which is 128 *\** 32 *\** 32 sthrough non-local block (Eq 2). Afterwards, to concate *H*^*′*^ with the output size of mid feature, bilinear interpolation is used for upsampling. Mid level features, *M R*^*C*H*W*^, from high resolution have a shape of 32×128x128, and this features go through a pointwise convolution layer and return *M*^*′*^ with same shape. Then, *M*^*′*^ is combined with context information with 1280 channels *g ∈ R*^*C*^. If we just simply concate *M*^*′*^ and context information, the representation of each pixel will be different due to the different size and resolutions of features, which may cause performance degradation in the segmentation task. To solve this problem, Adaptive Instance Normalization(AdaIN) proposed by [23] was used.

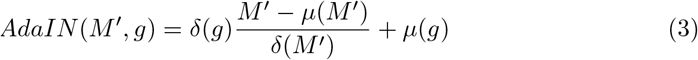

*AdaIN* receives input *M*^*′*^ as patch’s local information(content) from high resolution, and receives input *g* as patch’s global context information(style) from low resolution. *AdaIN* scales the *M*^*′*^’s normalized content input with *δ*(*g*), and shifts it with *μ*(*g*) without any learnable affine parameters (Eq 3). In short, *AdaIN* performs style transfer in the feature space by transferring feature statistics, specifically the channel-wise mean and variance.

**Figure 1.**
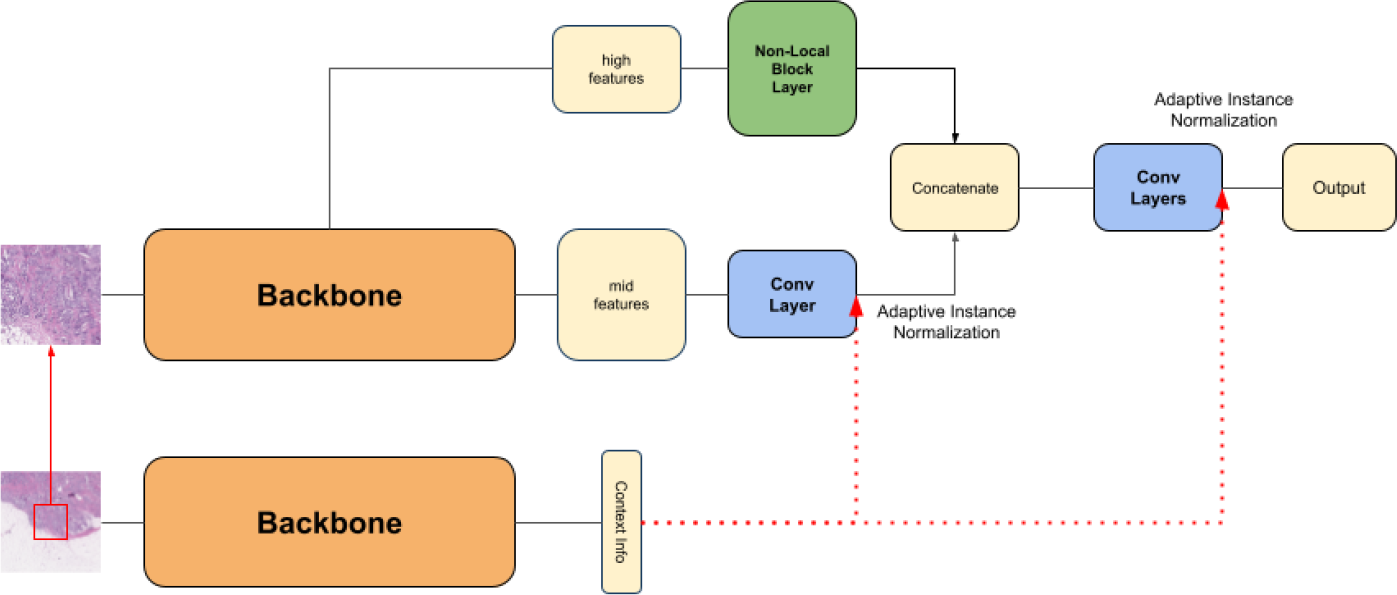
Multi-resolution Adaptive Normalization. When diagnosing a high resolution patch image (512×512 pixels, 50× magnitude), a model that simultaneously processes context information for the wide area in the low resolution patch image (512×512 pixels, 12.5x magnitude) using the adaptive normalization to calculate the final semantic segmentation.

**Figure 2.**
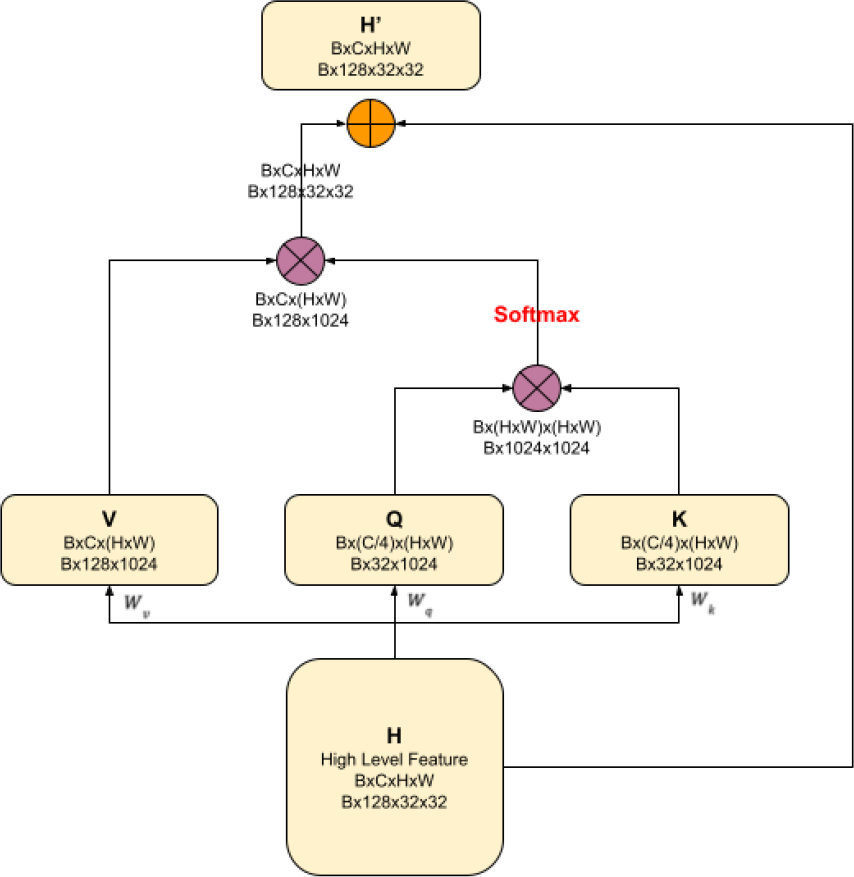
Non-Local Block. Query(*Q*), key(*K*), value(*V*) is constructed by pointwise convolution.

The patch-wise global feature from the Non-local block, *H*^*′*^, and *AdaIN* (*M, C*) combined through *AdaIN* are merged through a concatenation process. Afterward, we pass it through a convolutional layer, and after combining it with the context information through *AdaIN*, we obtain the final output.

### 3.2 Selective Segmentation method

In order to overcome the data uncertainties, we made a method ‘Selective Segmentation’ that evaluates uncertainty of regions. By using selective segmentation method, we aimed to address data uncertainty. [20] insisted that by using SelectiveNet, the model rejects those predictions it frequently gets wrong, which helped avoid uncertain predictions during both the inference and training processes. Inspired by SelectiveNet, we developed the selective segmentation method to apply to the segmentation task. The structure of the selective segmentation method follows the Figure 3.

**Figure 3.**
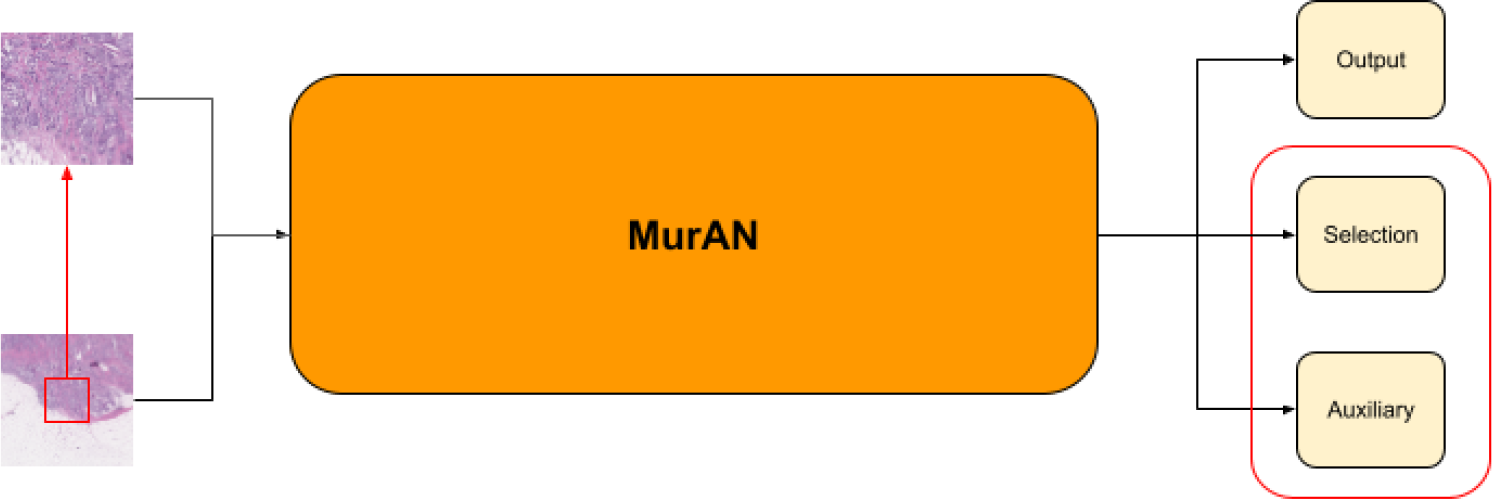
Selective segmentation method. Multi-resolution Selective Segmentaion model(MurSS) uses selective segmentation method. Selective segmentation method uses three heads. Original ouput head is to implement the prediction function *f* (*x*). Selection head is to implement the selection function *g*(*x*) to find the coverage. The auxiliary head is only used for training to generalize the loss.

Selective segmentation not only produces the output of MurAN but also produces two additional outputs: the selection output and the auxiliary output, resulting in a total of three distinct outputs. Among these, the function representing the original output is denoted as *f* (*x*), the function for the selection output as *g*(*x*), and the function for the auxiliary output as *h*(*x*). [20] states that the performance of selective method is quantified using coverage and risk. We defined the empirical coverage as

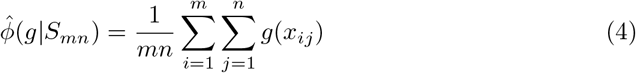

where *S*_*mn*_ is annotated data 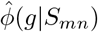shows the ratio of the region that is selected.Then, the empirical risk is defined as

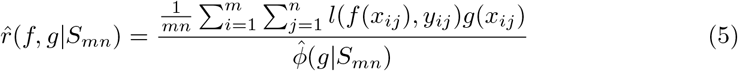

where, *l*(*f* (*x*_*ij*_), *y*_*ij*_) is cross entropy loss and

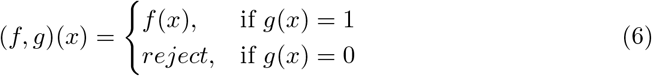

Also, quadratic penalty term Ψ(*a*) = *max*(0, *a*)^2^ is added as follow.

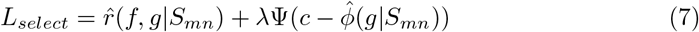

where *c* is the target coverage, and *λ* is controlling the relative importance of the constraints. We train MurSS to reduce the empirical selective risk in order to minimize *L*_*select*_. To find the optimal empirical selective risk, certain pixels must be rejected, under the condition that the percentage of non-rejected pixels meets or exceeds the coverage *c* that we have set. When 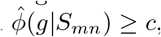quadratic penalty term will be 0 to reduce the *L*_*select*_. However, if we calculate the loss based on coverage region only, the model may be overfitted to the selected data. To address this, we use *h*(*x*), which represents the auxiliary prediction, to compute the auxiliary prediction risk during the training process. Therefore, our model’s loss is defined as

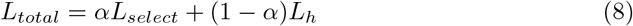

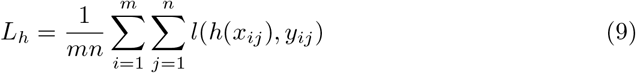

where *α* is hyperparameter to control the ratio of empiricial risk and auxiliary prediction risk, and *l*(*h*(*x*_*ij*_), *y*_*ij*_) is cross entropy loss.

## 4. Experiment

To measured the performance of MurSS and MurAN, we used Hematoxylin and Eosin(H&E) stained WSIs. These datasets are consisted of 1,181 TCGA Breast Invasive Carcinoma(BRCA) and 95 BRCA by Korea University Guro Hospital. To train MurSS and MurAN, three pathologists made annotations for benign, DCIS, and IDC, and 87 slides were excluded from the training phase due to slide quality issues. As a train dataset, 876 TCGA-BRCA WSIs were randomly selected and used, and 95 slides from Korea University Guro Hospital were used for validation. Finally, the remaining 218 TCGA-BRCA WSIs were used in the test set for the final performance comparison. We divided each slide into 512×512 pixel patches for training, evaluation, and testing. High resolution images were consisted of 512×512 pixels (50× magnitude), while low resolution images were consisted of 512×512 pixels (12.5× magnitude). When comparing the number of data based on the number of pixels from WSIs, there is an extreme imbalance for DCIS. Therefore, we oversampled the DCIS data by 9 times for training and performance comparison.

**Table 1.** Pixel-wise ratio for each class. Number of pixels is counted, and the ratio is calculated.

| Pixels ratio | Benign | DCIS | IDC |
| --- | --- | --- | --- |
| Base | 59.5% | 0.5% | 39.9% |
| After Oversampled | 57.3% | 4.3% | 38.4% |

### 4.1 Multi-resolution Adaptive Normalization model

To validate the performance of the MurAN, we compared it to various segmentation models such as U-Net [9], HRNet [11], and DeepLabV3 [10]. To compare the performance of each model, we compared pixel-wise accuracy and Intersection over Union (IoU) for benign, DCIS, and IDC. Table 2 shows the performance of each model. MurAN outperforms the other models, but all the models have a common problem of poor IoU for DCIS. However, MurAN shows better performance for DCIS than other models. When using the MurAN, the IoU of DCIS increased from a maximum of 11.4% to a minimum of 5.1%, and the mIoU of the entire classes increased from a maximum of 4.9% to a minimum of 2.7%.

**Table 2.** Performance table for basic segmentation models. Performance was measured by pixel-wise accuracy, mean Intersection of Union (mIoU) and Intersection of Union (IoU) for each class.

| Model | Accuracy | mIoU | IoU Benign | IoU DCIS | IoU IDC |
| --- | --- | --- | --- | --- | --- |
| U-Net | 0.948 | 0.724 | 0.930 | 0.370 | 0.868 |
| HRNet | 0.949 | 0.743 | 0.927 | 0.433 | 0.869 |
| DeepLabV3 | 0.944 | 0.721 | 0.922 | 0.385 | 0.856 |
| **MurAN** | **0.956** | **0.770** | **0.937** | **0.484** | **0.890** |

### 4.2 Multi-resolution Selective Segmentation model

There were limitations in achieving performance improvements for DCIS from results using MurAN. To overcome these performance barriers, various experiments were conducted, which identified the lack of data on DCIS and the morphological similarity between DCIS and IDC as the reason. To resolve the uncertainty of the data, we developed MurSS using MurAN with selective segmentation method. Also, following experiments were conducted.

- In order to reduce data uncertainty, regions with different annotation labels between pathologists and poor slide-quality patches were removed when training MurAN. Also DCIS data were added by reducing the stride when extracting DCIS patches from WSIs.
- In the process of calculating *ADAIN*, information may be damages and artifacts may occur which will make data more uncertain. Therefore, demodulation was applied which was proposed by [24].
- To overcome the uncertainty of data, selective segmentation method was applied to MurAN(MurSS). For the loss fuction, we used *λ* as 32, *c* as 0.95, and *α* as 0.5.

Table 3 shows the results of experiments conducted to resolve the data uncertainty that happened during the training phase of MurAN.

**Table 3.** Performance table for segmentation models to overcome data uncertainty. Performance was measured by pixel-wise accuracy, mIoU and IoU for each class.

| Model | Accuracy | mIoU | IoU Benign | IoU DCIS | IoU IDC |
| --- | --- | --- | --- | --- | --- |
| MurAN | 0.956 | 0.770 | 0.937 | 0.484 | 0.890 |
| MurAN + data | 0.960 | 0.789 | 0.942 | 0.527 | 0.899 |
| MurAN + demo | 0.961 | 0.792 | 0.943 | 0.532 | 0.901 |
| **MurSS** | **0.994** | **0.911** | **0.991** | **0.762** | **0.981** |

**Table 4.**
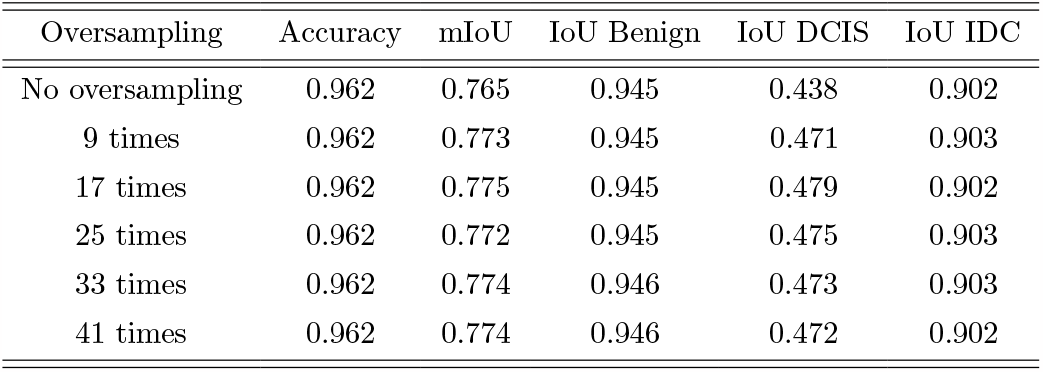
Performance table to overcome class imbalance through oversampling.

| Oversampling | Accuracy | mIoU | IoU Benign | IoU DCIS | IoU IDC |
| --- | --- | --- | --- | --- | --- |
| No oversampling | 0.962 | 0.765 | 0.945 | 0.438 | 0.902 |
| 9 times | 0.962 | 0.773 | 0.945 | 0.471 | 0.903 |
| 17 times | 0.962 | 0.775 | 0.945 | 0.479 | 0.902 |
| 25 times | 0.962 | 0.772 | 0.945 | 0.475 | 0.903 |
| 33 times | 0.962 | 0.774 | 0.946 | 0.473 | 0.903 |
| 41 times | 0.962 | 0.774 | 0.946 | 0.472 | 0.902 |

**Table 5.** Performance table to overcome class imbalance through weighted cross-entropy loss.

| Weight Ratio | Accuracy | mIoU | IoU Benign | IoU DCIS | IoU IDC |
| --- | --- | --- | --- | --- | --- |
| 1:1:1 | 0.962 | 0.773 | 0.945 | 0.471 | 0.903 |
| 1:3:3 | 0.957 | 0.766 | 0.938 | 0.465 | 0.894 |
| 1:9:9 | 0.947 | 0.747 | 0.922 | 0.444 | 0.873 |
| 1:13:13 | 0.943 | 0.738 | 0.915 | 0.434 | 0.915 |
| 1:3:1 | 0.961 | 0.773 | 0.944 | 0.476 | 0.900 |
| 1:9:1 | 0.961 | 0.765 | 0.945 | 0.448 | 0.902 |

We found that the IoU of DCIS increased by 4.3% when we reorganized the data and adjusted the stride to extracts more patches that contains DCIS. We also found that the IoU of DCIS increased by 4.8% when we added demodulation to MurAN’s *AdaIN* to reduce feature uncertainty. Finally, when using our proposed model MurSS, the IoU performance for DCIS increased by 27.8%. Also, the IoU of benign increased by 5.4%, and the IoU of IDC increased by 9.1%. It shows the total of 14.11% increasing rate in mIoU when using MurSS. Figure 4 shows the visualisation results from the annotation of the pathologist and model’s output. Since MurSS used selective segmentaion method, the model segment slides more accurately by understanding uncertainty of these regions.

**Figure 4.**
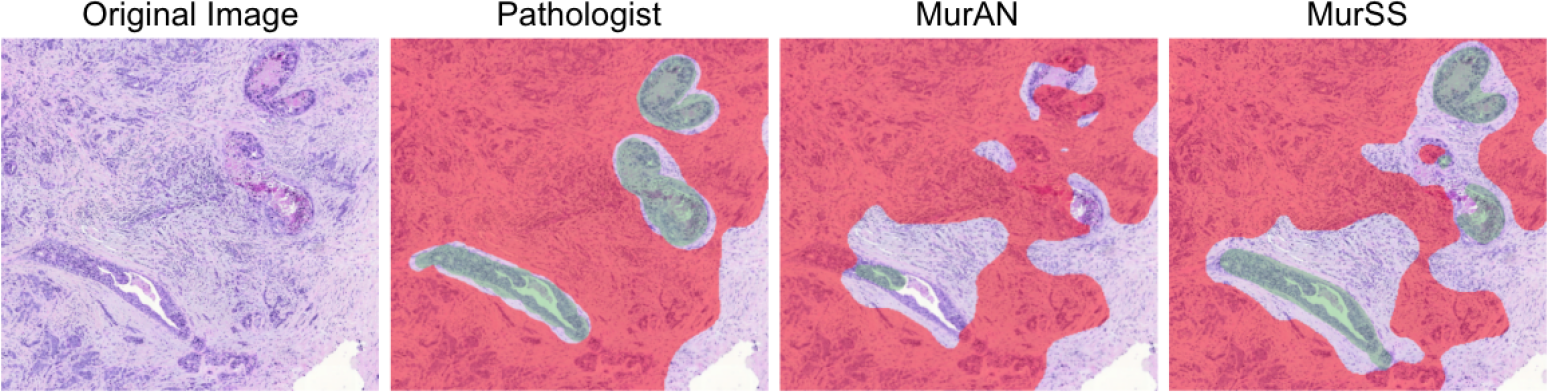
Annotation of the pathologist and models. With Selective Segmentation method, the model annotated slides more accurately by understanding uncertainty of these regions.

Figure 5 shows annotations of ambiguous regions according to predefined ratio *c* as 0.95, 0.90 and 0.80. Since pathologists usually annotate DCIS-like lesions as IDC if they are surrounded by IDC, these regions are ambiguous for models. We find that as the uncertainty ratio increased, the MurSS classified ambiguous regions as ‘uncertain’(purple). Using the MurSS, the performance of the model at regions labeled as certain hugely increased, which shows that the model distinguishes well between the certain and the uncertain regions.

**Figure 5.**
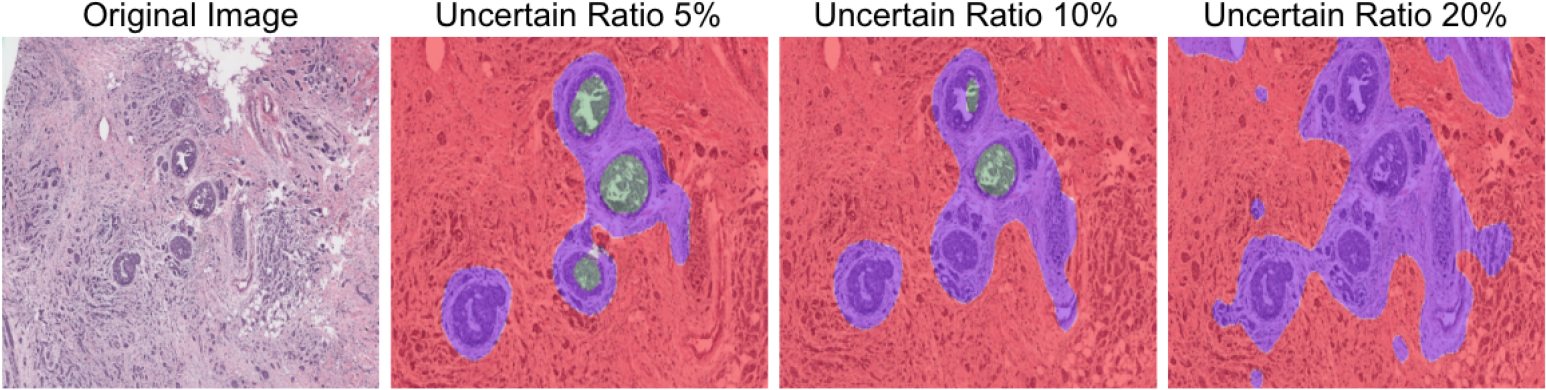
MurSS’s annotations of ambiguous regions according to predefined ratios 0.95, 0.90 and 0.80. MurSS classified ambiguous regions as ‘uncertain’(purple)

## 5. Conclusion

Due to the characteristics of medical image analysis, WSIs have to be divided into patches. It may focus on the local information rather than the whole slide images. Therefore, we proposed the Multi-resolution Adaptive Normalization model to use both high and low resolution images. Moreover, since medical image analysis can directly affect to patients, accurate diagnosis is important. However, the lack of well-labeled data and uncertain labeled data can make our model uncertain. Therefore, we suggest a method named ‘Selective Segmentation’ that automatically detects uncertain regions of WSIs. Through the Multi-resolution Selective Segmentation model, we could improve our model performance.

## Ablation Study

Various experiments were conducted to increase the IoU for DCIS and IDC of the MurAN. Since it takes a long time to train, 50 slides were randomly selected from the datasets and the experiment was organized. Oversampling and weighted sum cross-entropy loss were used.

